# Severe fever with thrombocytopenia syndrome virus and parthenogenetic Asian longhorned tick *Haemaphysalis longicornis* (Acari: Ixodidae)

**DOI:** 10.1101/2021.10.20.465214

**Authors:** Xing Zhang, Chaoyue Zhao, Chaoyuan Cheng, Hongyue Li, Tao Yu, Kevin Lawrence, Jimin Sun, Zeyu Yang, Ling Ye, Hongliang Chu, Ying Wang, Xiaohu Han, Yongchao Jia, Shuozhang Fan, Hirotaka Kanuka, Tetsuya Tanaka, Cheryl Jenkins, Kristene Gedye, Shona Chandra, Dana C. Price, Qiyong Liu, Young Ki Choi, Xiangjiang Zhan, Guogang Zhang, Zhibin Zhang, Aihua Zheng

## Abstract

Severe fever with thrombocytopenia syndrome virus (SFTSV) is spreading rapidly in Asia. It is transmitted by *Haemaphysalis longicornis* (Asian longhorned tick, ALT), which has both parthenogenetically and sexually reproducing populations. Parthenogenetic populations were found in at least 15 provinces in China and strongly correlated with the distribution of SFTS. The distribution of SFTS cases was however poorly correlated with the distribution of populations of bisexual ALTs. Phylogeographic analysis suggested that the parthenogenetic population spread much faster than bisexual population because colonization is independent of sexual reproduction. A higher proportion of parthenogenetic ALTs were collected from migratory birds captured at an SFTS endemic area, implicating the contribution to the long-range movement of ALTs in China. The SFTSV susceptibility of parthenogenetic females was like that of bisexual females under laboratory conditions. These results suggest that parthenogenetic ALT, probably transported by migratory birds, play a major role in the rapid spread of SFTSV.

**Article Summary Line:** The parthenogenetic population of Asian longhorned tick is broadly distributed in China and plays a major role in the long-distance spread of SFTSV and perhaps future invasion of countries outside of Asia.

## Introduction

Parthenogenesis is the development of an embryo from an unfertilized egg and is a common reproductive mechanism in invertebrate arthropods, especially insects and mites (1). One of the potential advantages of parthenogenesis is that the offspring are all genetically identical, and in relatively stable environments this can lead to a rapid expansion in numbers. Furthermore, following dispersal, new populations can be established from a single female. The Asian long-horned tick (*Haemaphysalis longicornis* Neumann, 1901; ALT) is one of a small number of medically important tick species that have both parthenogenetic (parth-) and bisexual (bi-) populations (2). The parth- population of ALT originated in Northern Japan (3) and is now common in the Asia-Pacific region. Both parth- and bi- populations occur in East Asia, but only the parth- population is found in Oceania (3). In China, the parth- population has only been reported in a few locations (4). In 2017, parth- ALT were found in New Jersey, USA (5) and by 2020, had been found across 12 states primarily within the Eastern USA (6).

Severe fever with thrombocytopenia syndrome virus (SFTSV) is a tick-borne phlebovirus transmitted by ALT and was first described in China in 2009 at the border of Henan, Anhui, and Hubei provinces (7, 8). SFTSV is maintained and transmitted by ALTs in the larva, nymph, and adult stages in both transovarial and transstadial modes (8-10). Human mortality rates resulting from SFTSV infection range from 6% to 30% (7, 11). From 2011 to 2016, SFTS cases were reported in 18 of the 34 provinces in China, with a 3-fold increase in the case number (from 500 cases to 1500 cases) (11). Most cases were in the rural areas of Henan (37%), Shandong (26.6%), Anhui (14%), and Hubei (12.6%) (11). SFTSV has also been reported in Republic of Korea (2012), Japan (2014), Vietnam (2019), and Pakistan (2020) (12-15). Phylogenetic analysis revealed that SFTSV isolates separate into the Chinese clade and the Japanese clade, which is consistent with their geographic distribution (16). A close relative of SFTSV, Heartland virus, was reported in the USA in 2012 transmitted by *Amblyomma americanum* (17, 18).

ALT is the dominant tick species in the SFTSV endemic area, with proportional representation in the tick fauna of 88% in Jiangsu Province, China, and 91% in Gangwon Province, Republic of Korea (19, 20). Most of the SFTSV endemic areas are rural, and 97% of the patients are farmers living in wooded and hilly areas, far from modern transportation and cities (21). The rapid spread of SFTSV is currently unexplained, although ALT has a broad host range (18), enabling several possible modes of dissemination. For example, livestock and wild mammals are common hosts for ALT so the grazing movement of cattle or foraging of wild mammals, such as hares, could rapidly distribute ticks in an area with a suitable habitat (22). Furthermore, birds range far enough daily to transport ticks within a district, while long-range dispersal of tick-borne pathogens can also be accelerated by tick-infested migratory birds (23). In 2016, SFTSV antibodies were detected in tick-infested migratory Greater White-fronted Geese in Jiangsu Province, China (24). This finding has led to the suggestion that migratory birds may have been involved in the rapid spread of this disease within China. Two areas of China that have high endemic levels of SFTSV are both situated on major bird migratory routes (25). The Dabie Mountain area region (where SFTSV first reported), is in the middle of an important bird migration route from Dongting Lake and Poyang Lake to Siberia, while Penglai City and Dalian City are both located on the northern part of the Asia-Pacific migratory route. Dongting Lake and Poyang Lake are two of the most important overwintering sites for migratory birds in China (26, 27).

The aim of this study was to test the hypothesis that parth- ALT, possibly carried by migratory birds, are responsible for the rapid spread of SFTSV in China and Asia. To test this hypothesis we carried out a series of linked experiments, which included mapping the distribution of bi- and parth-ALTs in those Chinese provinces with a high prevalence of SFTSV, estimating the geographical correlation between bi- and parth- ticks and SFTSV cases for those provinces, surveying the infestation of ALT in migratory birds in a high prevalence SFTSV city, testing the virus acquisition and transstadial passage of ALTs for SFTSV, inferring the phylogeny of ALTs using ticks collected from around the world, and estimating the correlation between migratory bird routes and bi- and parth- ticks populations within China.

## Materials and Methods

### Tick Collection in China

ALTs were collected from 73 counties covering 20 provinces where SFTSV is endemic (Fig. 1A and S2). Ticks of all life stages were collected by flag-dragging and removal directly from animals between April and November 2019. Ticks were identified based on morphological characteristics, visualized through a light microscope, with further molecular confirmation in the laboratory by sequencing the mitochondrial 16S ribosomal RNA (16S rRNA) gene. Primers: (16S-1) CTGCTCAATGATTTTTTAAATTGCTGTGG and (16S-2) CGCTGTTATCCCTAGAGTATT. A single leg was removed from each tick for the molecular analysis to confirm identification. A random sample of 5 or 6 live tick from each county sampled were then stored at room temperature until ploidy detection, whilst the remaining tick specimens were stored at -80°C.

**Figure 1.**
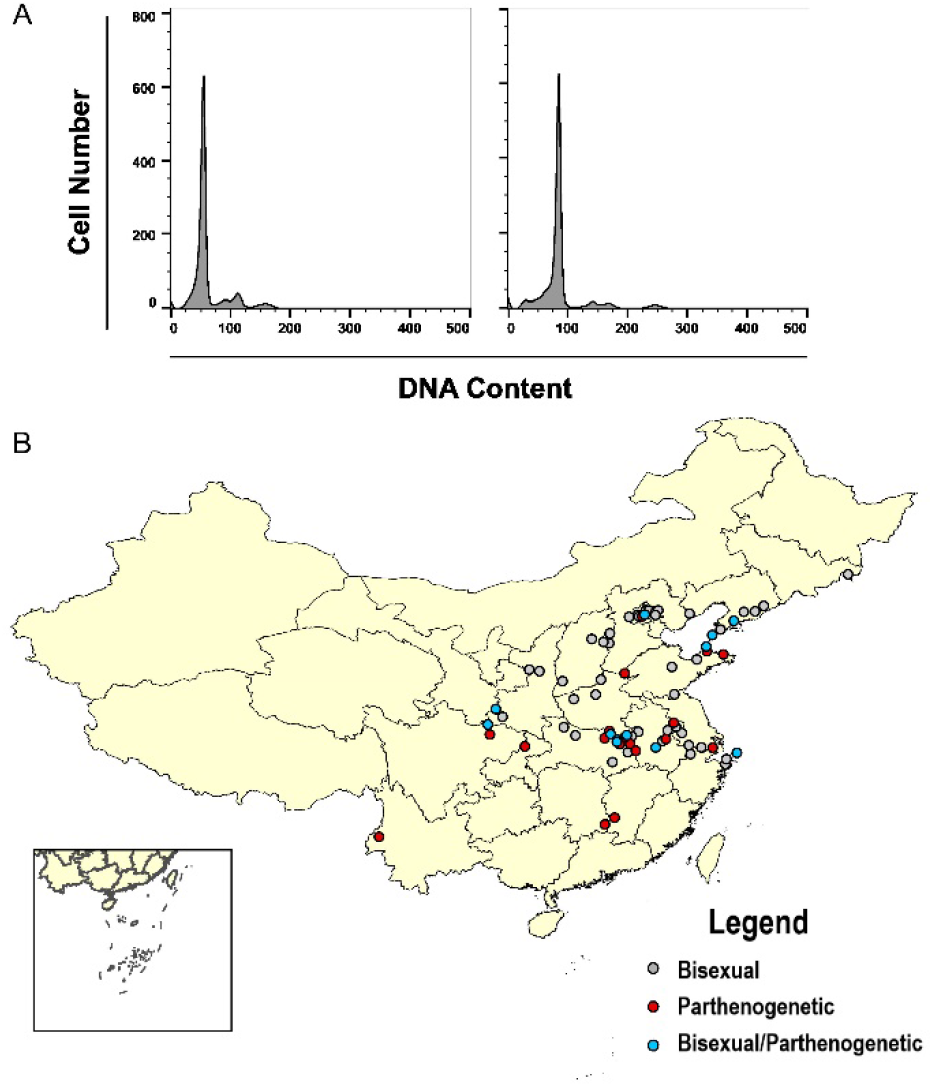
Polyploid analysis and geographical distribution of bi- and parth- Asian longhorned tick (ALT) populations. (A) The ploidy of ticks ALTs was tested using flow cytometry by measuring the fluorescence intensity of cell nucleus stained with DAPI. Illustrated is a representative of bi-(2n) with peak at the 66 position and parth-(3n) at 99. (B) Geographical distribution of bisexual and parthenogenetic ALTs collected in China. Red dots indicate parth-; gray dots indicate bi-; blue dots indicate both bi- and parth-.

### Tick collection overseas

Eight extracted tick DNA samples were collected from overseas collaborators including Japan, Republic of Korea, New Zealand, Australia, and the USA in 2019 and subjected to the same molecular analysis as the Chinese collected ALT ticks.

### Polyploid analysis of ticks

Bi- and parth- ticks are difficult to distinguish using classical taxonomic methods, but they can be identified by karyotype analysis, so flow cytometry was used to test the ploidy of the tick chromosomes (4). This was accomplished by measuring the fluorescence intensity of cell nuclei stained with DAPI (4’ ,6-diamidino-2-phenylindole). Sysmex Partec CyFlow Space (Sysmex Partec, Görlitz, Germany) was used in this analysis and Sysmex Partec CyStain DNA 1 Step kit was used.

### Correlation between bi- and parth- ALTs and SFTS cases

The geographical correlation between SFTS cases and the distribution of different populations of ALT was analyzed by linear regression (28). The total number of recovered ticks, aggregated at the province or municipality level, was used as the independent variable, and the incidence of SFTS cases (cases per million population) reported in each respective province or municipality was used as the dependent variable. SFTS cases data summarized by province or municipality for the year 2019, were obtained from the Chinese Center for Disease Control and Prevention.

### Phylogenetic tree and genetic distance

For each county, one bi- tick sample or/and one parth- tick sample were picked randomly for whole mitochondrial sequencing. The whole mitochondrial genomes were sequenced by next-generation sequencing (Tsingke, Beijing, China). Phylogenetic analysis was performed using the whole mitochondrial genomes of 46 bi- and 35 parth- ticks collected from China, Japan, Republic of Korea, New Zealand, Australia, and the USA. Tick DNA was extracted using the MightyPrep reagent for DNA Kit (Takara, Japan) according to the manufacturer’s instructions. The mitochondrial DNA were sequenced by next generation sequencing (Tsingke Biotech, Beijing, China) and deposited in GenBank (Accession number MW642336-MW642407). Two phylogenetic trees were constructed using this data, the first using the maximum likelihood method, MEGA-X with the bootstrap value set at 1000 and the second using the Bayesian interference method, MrBays-3.2.7 (http://nbisweden.github.io/MrBayes/index.html) performing 1,500,000 generations. The genetic distance (GD) used to calculate the dispersal index was equal to the nucleotide substitution rate.

### Nucleotide diversity

The nucleotide diversity (Pi) in each dataset is estimated by:

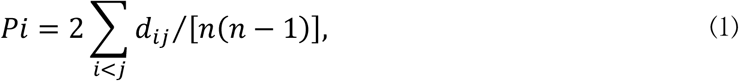

where *d_ij_* is an estimate of the number of nucleotide substitutions per site between DNA sequences i and j and n is the number of DNA sequences examined (29). The calculation for Pi was implemented in DnaSP-V6 (30).

### Dispersal index

Without longitudinal surveillance data, we could not evaluate the spread velocity of ticks directly, so we defined a dispersal index to quantify and compare the dispersal ability of bi- and parth- ticks. According to the molecular clock theory, the mutation rate is relatively constant, so genetic distance can be used to represent time distance (31). The dispersal index (I) was defined as: I=D/G. Where, D represents the sum of the spatial distances between every two samples (i.e. pairwise geographical distance), and G represents the genetic distance (nucleotide substitutions) between every two samples (i.e. pairwise genetic distance). This index is not the true dispersal velocity itself, but we can infer that the greater the dispersal index value then the greater the dispersal velocity must be. An independent sample t-test was performed to contrast the dispersal rates for bi- and parth- ticks. The calculation of dispersal index and t-test were implemented by customed scripts written in Python-3.7 (https://www.python.org).

### Migratory bird capture and tick collection

To test our hypothesis that migratory birds are carrying ALT ticks, we investigated the presence of ticks on northward migratory birds at a high-density stopover site at Penglai (37°55′13.84″N, 120°43′53.27″E), Yantai City, Shandong Province in April 2021. Shandong Province is an important location on the Asia-Pacific migratory route. Birds were captured with mist nets (2.5 × 6 m or 2.5 × 12 m, mesh size:3.0 cm) placed in a wooded habitat. Upon capture, birds were meticulously examined for the presence of ticks by searching the ear canals, back of the head, mandibular area, and perimeter of the eyes. Ticks were then removed with fine forceps.

### Penglai City tick diversity

Penglai City is one of the most endemic areas of SFTS in China and an important location in the Asia-Pacific migratory route. Parth- ticks were collected and sequenced from nine locations in Penglai City. Phylogenetic analysis was used to compare the diversity of ticks from Penglai city to 15 other provinces in China.

### Virus acquisition and transstadial passage of ALTs for SFTSV

The virus susceptibility of parth- and bi- populations for transmitting SFTSV was tested using laboratory-adapted ALT colonies and an interferon α/β receptor knockout (IFNAR^−/−^) mouse model. Nymphal ALTs were infected by feeding on IFNAR^−/−^ C57/BL6 mice, previously inoculated with 2 × 10^3^ FFU of SFTSV (Wuhan strain; GenBank accession numbers: S, KU361341.1; M, KU361342.1; L, KU361343.1). The fed nymphs were collected after they were fully engorged and had detached from the mice. SFTSV RNA levels in the ticks were analyzed after they molted into adults. Total RNA prepared from the homogenates of the ticks was extracted using TRIzol reagent (Thermo Fisher Scientific, USA) according to manufacturer instructions. Samples were analyzed using a One-Step SYBR PrimerScript reverse transcription (RT)-PCR kit (TaKaRa, Japan) on an Applied Biosystems QuantStudio. Each sample was measured in triplicate. The primers were designed as previously described [32].

## Results

### Tick distribution and ploidy analysis

There were 1328 ALTs confirmed by 16S rRNA sequencing, of which 271 (20.4%) live ticks were further submitted for ploidy analysis by flow cytometry (255 ticks) or by mitochondrial sequencing (16 ticks) (Table S1). Ploidy testing showed a peak for bi- (diploid) population at the 66 position and for the parth- (triploid) population at the 99 position (Fig. 1A). Of the ticks submitted for ploidy analysis, 186/271 (69%) were identified as bi- and 85/271 (31%) were identified as parth-. Bi- ticks were detected in 55/73 (75%) of counties, parth- ticks were detected in 30/73 (41%) of counties, and a mixture of both populations were detected in 12/73 (16%) counties (Fig. 1B and Table S1). Interestingly in 18/73 (25%) of counties only parth- ticks were found and in 43/73 (59%) of counties only bi- ticks were found.

### Correlation of SFTSV with bi- and parth- populations of ALTs

SFTSV cases showed a strong correlation with the parth- population (*R*^*2*^ = 0.685, *p* < 0.001), but almost no correlation with the bi- population (*R*^*2*^ = 0.026, *p* = 0.501) (Fig. 2). In the highly endemic Dabie mountain area (located at the border of Henan, Anhui, and Hubei provinces in central China), 11 out of 14 counties 66% of the collected ALTs were parth- (Table S1). These results suggest that the parth- populations of ALT are geographically strongly associated with cases of SFTSV.

**Figure 2.**
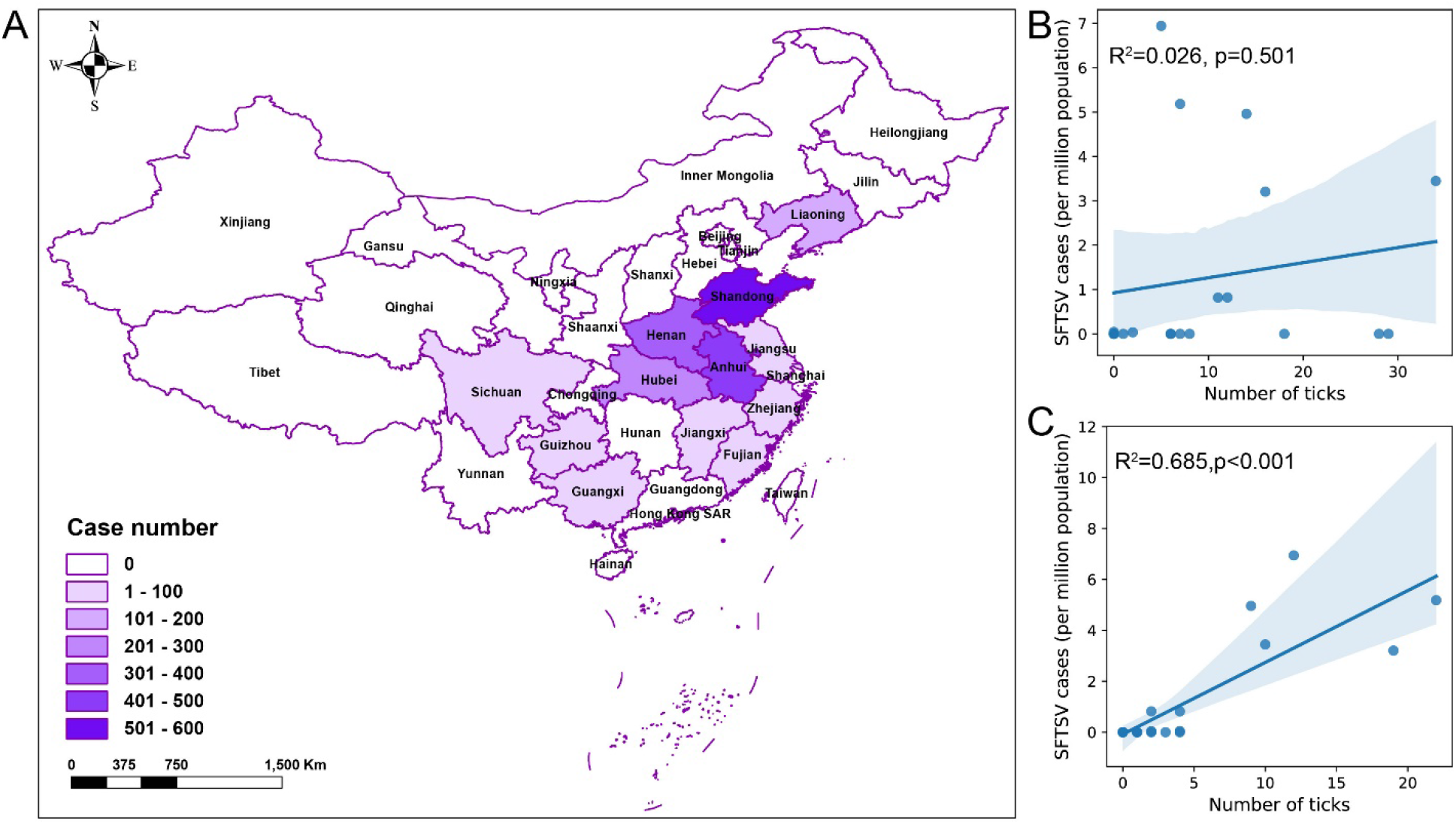
The SFTS endemic area is highly correlated with the parth- Asian longhorned tick (ALT) population. (A) The distribution of Severe fever with thrombocytopenia syndrome (SFTS) cases in China in 2019 (China CDC). B and C show that the correlation between incidence of SFTS cases (cases per million population) and the number of bi- population (B) and the parth-population (C) ALTs in different provinces. Each dot represents a province.

### Phylogenetic analysis of bi- and parth- populations

For each county, one bi- or/and parth- ALTs were submitted for mitochondrial sequencing. Eighty one whole mitochondrial genomes were obtained from 73 Chinese ticks and 8 ticks outside of China. Fig. 3A, Fig. S3 and Fig. S4 clearly show that the parth- and bi- populations divide into two distinct lineages that can be discriminated by a single T deletion at nucleotide 8497 in the un-translational region (Fig. S5). This suggests that the parth- population may have originated from a single event without gene exchange. The mean genetic distance (GD) between all sequences was 0.0078 as measured by nucleotide substitution rate. The New Zealand and Australian parth-strains were very similar to the parth-strain from Okayama, Japan (Mean GD = 0.0003). The parth-strain from Kagoshima, Japan, was a close relative to strains collected from Beijing, Hubei, and Henan (Fig. S1A ⑧④②), which are geographically separate. The strain from New Jersey, USA, was very similar to the strain from Jeju Island, Republic of Korea (GD = 0.0001).

**Figure 3.**
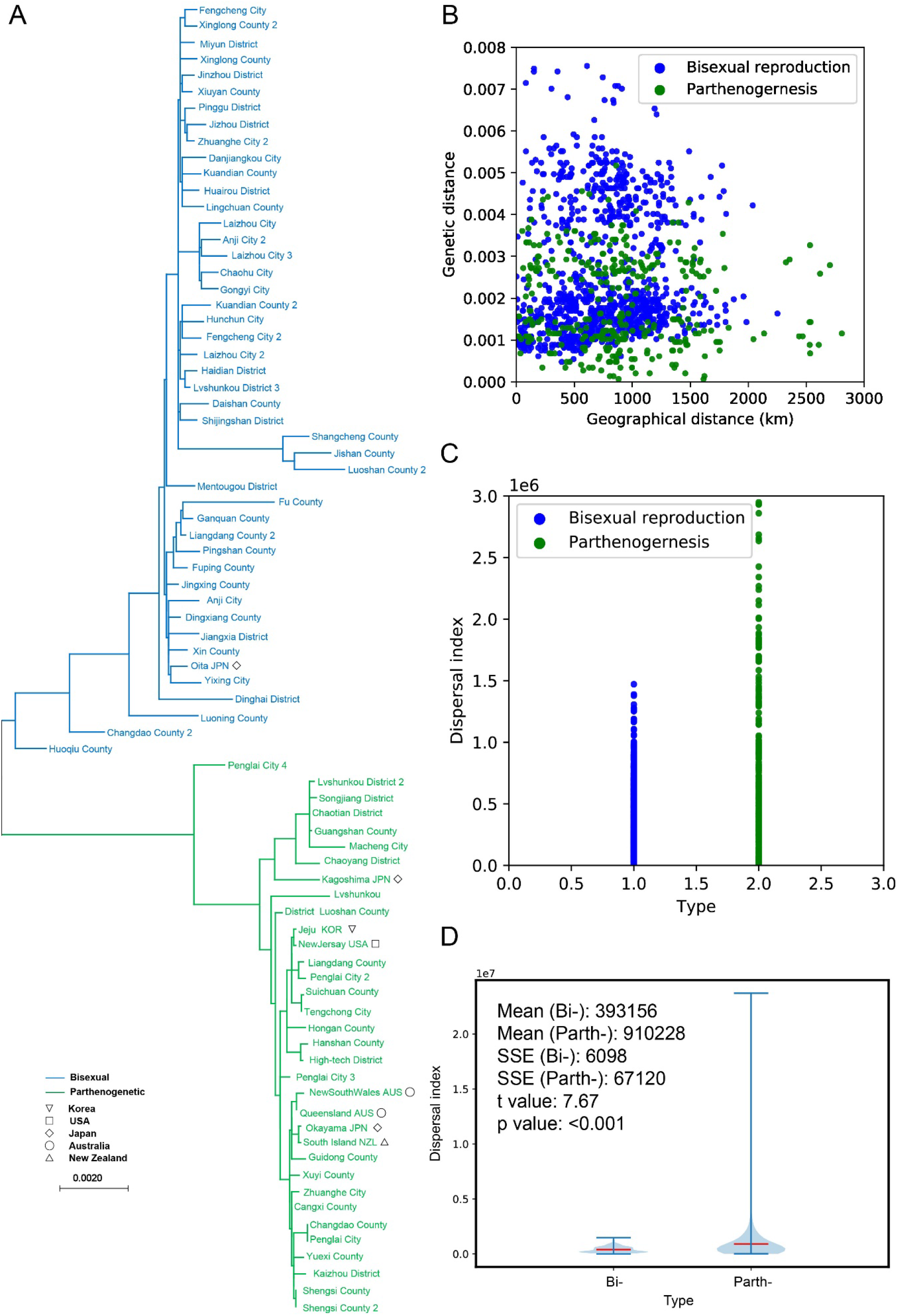
Phylogenetic and phylogeographic analysis of the bi- and parth- Asian longhorned ticks (ALTs). (A) Maximum likelihood tree established with the mitochondrial genomes of ALTs collected in the Asian-Pacific region. Multiple ALTs from the same county were marked by number. (B) Distribution of bi- and parth- ALTs in pairwise genetic distance and pairwise geographical distance. (C) Distribution and (D) the differences of dispersal index (see methods) between bi- and parth- ALTs. The red bar in the violin plot represents the mean dispersal index, and the shaded blue areas indicate the kernel density estimation of the dispersal indices.

### Genetic diversity

Despite the loss of sexual reproduction, high genetic diversity has been reported in the asexual populations of many different species (32). The genetic diversity (Pi) of the bi- and parth- population, as measured by the mitochondrial genome, was 0.00249 and 0.00188, respectively. These results indicate that the genetic diversity of the bi- and parth- population was similar and that the parth- population may have diverged from the bi- population at a very early age.

### Dispersal index of bi- and parth- ticks

Compared with bi- ticks, parth- ticks have a wider pairwise geographical distance distribution and a narrower pairwise genetic distance distribution (Fig. 3B). The dispersal index of parth- ticks was significantly higher than bi- (t = 7.67, p < 0.001), with the mean dispersal index for parth- ticks (910228) being 2.3 times higher than for bi- ticks (393156) (Fig. 3C, D), indicating that parth- ticks have a higher dispersal capacity.

### The correlation between migratory birds and ALTs

Migratory birds were collected and examined for ALTs at Penglai City, which is a high SFTSV endemic area and located in the Asia-Pacific migratory route (Fig.S1 ⑩). Ninety-five birds belonging to 17 species were netted however, ALTs were only found on four species (n=54); Naumann’s thrush (*Turdus naumanni)*, the Grey-backed Thrush (*Turdus hortulorum)*, the great tit *(Parus major)*, and the chestnut-eared bunting (*Emberiza fucata)*. Only a total of 27 ticks were recovered from these birds of which 19/27 (70%) were identified as ALT, with 17/19 (89%) ALTs being parth-(Table 1). All recovered ALTs were nymphs.

**Table 1.** *Haemaphysalis longicornis* tick (ALT) collected from migratory birds and their hosts in Penglai City 2021.

| Avian host | No. of examined<br>birds | No. of birds with<br>ticks | No. of<br>ticks | No. of<br>ALT | Percent of Parth-<br>(%) |
| --- | --- | --- | --- | --- | --- |
| *Turdus*<br>*naumanni* | 45 | 8 | 11 | 3 | 33 |
| *Turdus*<br>*hortulorum* | 7 | 2 | 8 | 8 | 100 |
| *Parus major* | 1 | 1 | 5 | 5 | 100 |
| *Emberiza fucata* | 1 | 1 | 3 | 3 | 100 |

### Penglai city tick diversity

Phylogenetic analysis showed that the mitochondrial sequences of the parth- ALTs collected in Penglai city from vegetation were highly diverse when aligned with those from 15 provinces in China (Fig. S6). These data suggest that ticks of many different provinces were present in Penglai city and likely spread there by migratory birds.

### Virus acquisition and transstadial passage of ALTs for spreading SFTSV

A robust viremia was detected in the mice inoculated with SFTSV (Fig. 4A). After feeding until engorgement and moulting, both the parth- and bi- population developed average titers of 3 log RNA copies/mg without obvious differences (Fig. 4B). The SFTSV-acquisition and transstadial passage efficiency of the parth- population appeared comparable to that of the bi- population.

**Figure 4.**
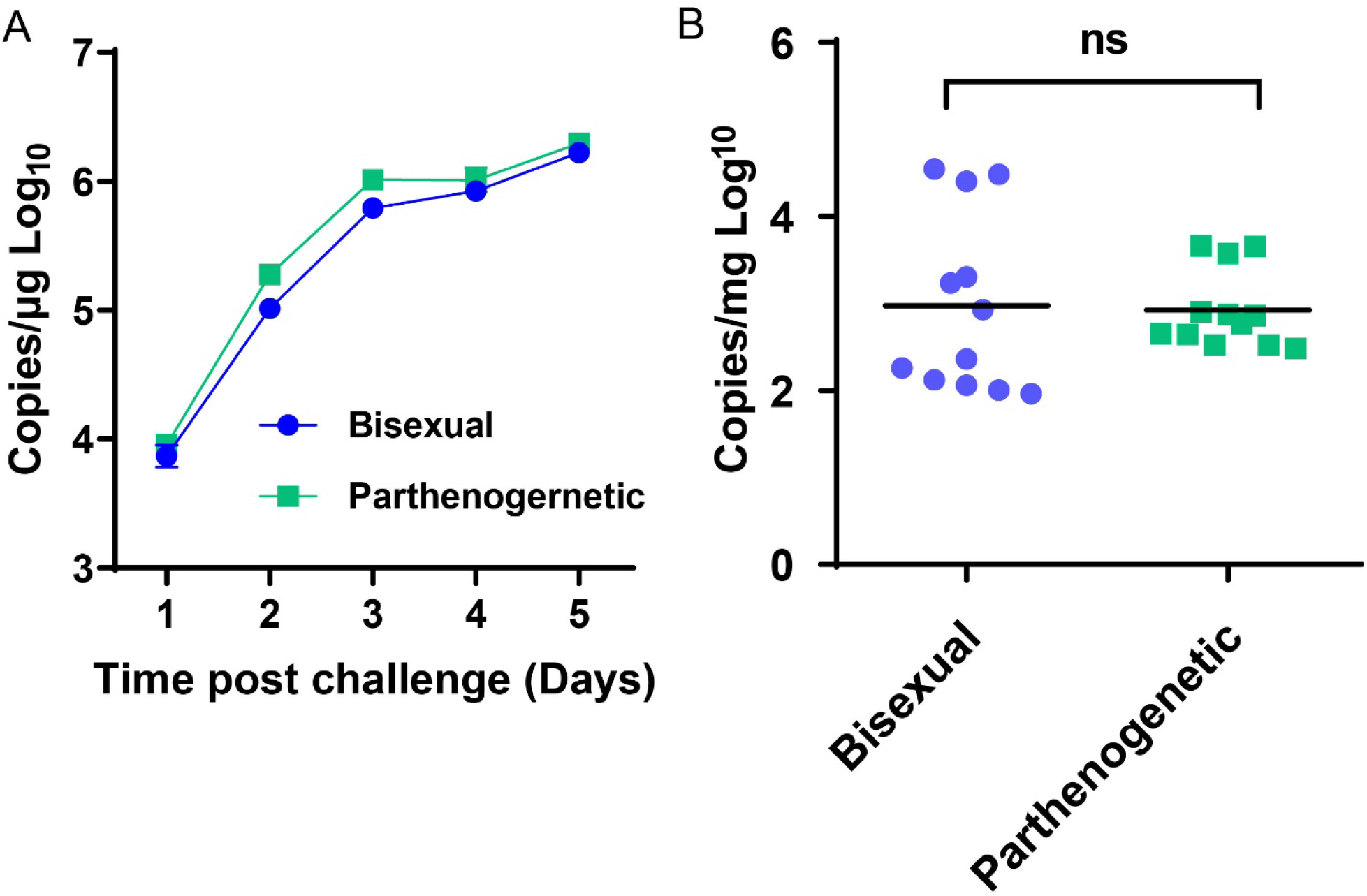
Susceptibility of bi- and parth- Asian longhorned ticks (ALTs) to Severe fever with thrombocytopenia syndrome virus (SFTSV). Groups of bi- or parth-nymph ALTs were fed separately on one IFNAR^-/-^ C57/BL6 mouse that was intraperitoneal inoculated with 2×10^3^ FFU of SFTSV. (A) Viremia of IFNAR^-/-^ C57/BL6 mice were monitored by Real-time PCR during tick feeding. (B) SFTSV infection in the ALTs were tested by Real-time PCR after molting into adult. Each dot represents one tick.

## Discussion

We found that the parth- population of ALT is more widely distributed in China than previously thought and that the distribution is highly correlated with the endemic regions of SFTSV. Phylogeographic analysis suggests that the parth- ALT population has spread more rapidly over a greater distance than the bi- population and assessment of virus acquisition and transstadial passage showed that bi- and parth- populations were comparable in maintaining the local transmission of SFTSV. Although only a small number of ticks were recovered, parth- ALTs were found dominant in migratory birds collected at a SFTS endemic area. We suggest that together these results strongly support the hypothesis that parth- ALT populations are responsible for the rapid spread of SFTSV within China, most likely through being disseminated by migratory birds.

If, as we suggest, migratory birds have played an important role in the spread of parth- ALT then this would partly explain the wide distribution of these ticks from the cold Far East of Russia to the tropical areas of Australia and the Fiji Islands. However, the important role of livestock, wild mammals, and humans via companion animals in the translocation of parth- ALTs should not be overlooked (22).

Migratory birds have long been known to be carriers of ticks. Penglai City is one of the most endemic areas of SFTS and is a key passage in the northern part of East Asian-Australasian Flyway. In this area, 96% of ALTs were parth- and showed extremely high diversity (Fig. S6). During a spring bird survey in Penglai City in 2021, ALTs were specifically found in four bird species *Turdus naumanni, Turdus hortulorum, Parus major*, and *Emberiza fucata* and 89% of them were parth-. Among the four bird species, *Turdus naumanni, Turdus hortulorum*, and *Emberiza fucata* migrate between East Asia and Siberia, and are occasionally found in Alaska (www.ebird.org). The preferred habitats for these 4 species are grasslands and bushes, which are also the preferred habitats of ALTs. These results suggest that migratory birds have an important role in the long-range movement of parth- ticks within China and potentially even trans-oceanic spread of SFTSV.

Parth- ALTs are also implicated in the spread of a pathogenic form of the blood parasite, *Theileria orientalis* throughout the Asia-Pacific (18). ALTs are purported to have been introduced to Australia in the 19^th^ century from Northern Japan and later disseminated to New Zealand, New Caledonia, and Fiji. This theory is supported by phylogenetic results of this study which show that the New Zealand and Australian ALTs are alike and closely resemble the parth- strain from Okayama, Japan (33). *T. orientalis* has been present in Australia for more than 100 years, having been introduced with the vector tick, and until 2006 caused only minor symptoms in livestock (34). In 2006, the pathogenic Ikeda genotype of *T. orientalis* was introduced from Eastern Asia into New South Wales, Australia (35) and by 2014, had spread to most of the states (36). The recent spread of *T. orientalis* across the Asia-Pacific and into North America highlights the risk of rapid disease agent transmission into areas where a competent vector (ALT) is already established. Thus, while SFTSV has not yet been detected in the Western Hemisphere, the presence of ALT parth- ticks in several countries within this region presents a clear risk for the emergence of this disease in the future.

### Ethics statement

All animal studies were carried out in accordance with the recommendations in the Guide for the Care and Use of Laboratory Animals of the Ministry of Science and Technology of the People’s Republic of China. The protocols for animal studies were approved by the Committee on the Ethics of Animal Experiments of the Institute of Zoology, Chinese Academy of Sciences (Approval number: IOZ-IACUC-2020-062).

## Acknowledgments

We thank Dr. Chaodong Zhu from Institute of Zoology, CAS for a critical review of this MS. We thank Dr. Shu Shen (Wuhan Institute of Virology, CAS), Dr. Jingwen Wang (Fudan University, Shanghai, China) for sending us ALTs. This project was supported by the National Science and Technology Major Project (2018ZX10101004); State Key Research Development Program of China (2019YFC12005004); National Natural Science Foundation of China, General Program (81871687) and Open Research Fund Program of State Key Laboratory of Integrated Pest Management (IPM1603 and IPM1806).

## Author bio

Dr. Zhang is an associate professor at College of Life Sciences, University of Chinese Academy of Sciences, China. His research interest includes vector biology and genetics, particularly in vector-borne pathogens and endosymbionts in ticks.

