## Supplemental figures for "Severe fever with thrombocytopenia syndrome virus and parthenogenetic Asian longhorned tick *Haemaphysalis longicornis* (Acari: Ixodidae)"

**Appendix information**

**Appendix Table.**

**Table S1.** Location and polyploid information of Asian longhorned ticks (ALTs) samples in China.

| | Location | Identified ticks | Identified ALT | Bi-ALT | Parth-ALT |
|---|---|---|---|---|---|
| # | Yuexi County, Anqing City, Anhui Province | 3 | 3 | | 3 |
| | Nanqiao District, Chuzhou City, Anhui Province | 24 | 1 | 1 | |
| | Chaohu City, Hefei City, Anhui Province | 2 | 2 | 2 | |
| # | Lujiang County, Hefei City, Anhui Province* | 10 | 7 | 1 | 6 |
| # | Huoqiu County, Lu 'an City, Anhui Province | 1 | 1 | 1 | |
| # | Jinzhai County, Lu 'an City, Anhui Province | 14 | 13 | | 1 |
| | Hanshan County, Maanshan City, Anhui Province | 5 | 5 | | 2 |
| | Chaoyang District, Beijing | 20 | 13 | | 3 |
| | Haidian District, Beijing | 60 | 60 | 6 | |
| | Huairou District, Beijing | 8 | 8 | 2 | |
| | Mentougou District, Beijing | 50 | 50 | 2 | |
| | Miyun District, Beijing | 30 | 30 | 6 | |
| | Pinggu District, Beijing | 29 | 29 | 6 | |
| | Shijingshan District, Beijing | 50 | 50 | 6 | |
| | Shunyi District, Beijing | 27 | 27 | 5 | 1 |
| | Liangdang County, Longnan City, Gansu Province | 20 | 20 | 7 | 1 |

| | | | | | |
|---|---|---|---|---|---|
| | Fuping County, Baoding City, Hebei Province | 34 | 33 | 6 | |
| | Xinglong County, Chengde City, Hebei Province | 20 | 10 | 3 | |
| | Haigang District ,Qinhuangdao City, Hebei Province | 18 | 9 | 2 | |
| | Jingxing County,Shijiazhuang City,Hebei Province | 20 | 20 | 6 | |
| | Pingshan County, Shijiazhuang City, Hebei Province | 38 | 30 | 5 | |
| | Luoning County, Luoyang City, Henan Province | 34 | 10 | 1 | |
| | Nanle County, Puyang City, Henan Province | 21 | 20 | | 1 |
| # | Gushi County, Xinyang City, Henan Province | 5 | 5 | 1 | |
| # | Guangshan County, Xinyang City, Henan Province | 3 | 3 | | 3 |
| # | Luoshan County, Xinyang City, Henan Province | 56 | 56 | 4 | 2 |
| # | Pingqiao District, Xinyang City, Henan Province | 8 | 8 | | 6 |
| # | Shangcheng County, Xinyang City, Henan Province | 20 | 11 | 2 | 1 |
| # | Xinxian County, Xinyang City, Henan Province | 14 | 6 | | 6 |
| | Gongyi City, Zhengzhou City, Henan Province | 7 | 7 | 2 | |
| # | Hongan County, Huanggang City, Hubei Province | 51 | 40 | 2 | 1 |
| # | Luotian County, Huanggang City, Hubei Province | 8 | 6 | 5 | |
| # | Macheng City, Huanggang City, Hubei Province | 6 | 6 | | 6 |
| | Danjiangkou City, Shiyan City, Hubei Province | 11 | 1 | 1 | |
| # | Guangshui City, Suizhou City, Hubei Province | 20 | 20 | | 2 |
| | Jiangxia District, Wuhan City, Hubei Province | 13 | 10 | 6 | |
| | Nanzhang County,Xiangyang City, Hubei Province | 9 | 9 | 1 | |
| | Guidong County, Chenzhou City, Hunan Province | 2 | 1 | | 1 |
| | Hunchun City, Yanbian Prefecture, Jilin Province | 50 | 50 | 6 | |

| | | | | |
|---|---|---|---|---|
| Xuyi County, Huai'an City, Jiangsu Province | 86 | 50 | | 4 |
| Donghai County, Lianyungang City, Jiangsu Province | 4 | 1 | 1 | |
| Jiangning District, Nanjing city, Jiangsu Province | 12 | 12 | 1 | |
| Liuhe District, Nanjing city, Jiangsu Province | 12 | 11 | 1 | |
| Wuzhong District, Suzhou city, Jiangsu Province | 8 | 8 | 1 | |
| Yixing City, Wuxi City, Jiangsu Province | 12 | 10 | 6 | |
| Suichuan County, Ji 'an City, Jiangxi Province | 21 | 21 | | 4 |
| Xiuyan County anshan City Liaoning Province | 14 | 10 | 5 | |
| Jinzhou District, Dalian City, Liaoning Province | 32 | 22 | 6 | |
| Lvshunkou District, Dalian City, Liaoning Province | 12 | 8 | 3 | 3 |
| Zhuanghe City, Dalian City, Liaoning Province | 13 | 12 | 1 | 5 |
| Fengcheng City, Dandong City, Liaoning Province | 27 | 27 | 6 | |
| Kuandian County, Dandong City, Liaoning Province | 30 | 10 | 3 | |
| Penglai City, Yantai City,Shandong Province | 31 | 10 | | 3 |
| Changdao County, Yantai City,Shandong Province* | 16 | 16 | 1 | 2 |
| High-tech District, Weihai City, Shandong Province | 34 | 34 | | 6 |
| Qingzhou City, Weifang City, Shandong Province | 50 | 50 | 1 | 2 |
| Laizhou City, Yantai City, Shandong Province | 3 | 1 | 1 | |
| Lingchuan County, Jincheng City, Shanxi Province | 27 | 16 | 2 | |
| Dingxiang County, Xinzhou City, Shanxi Province | 9 | 9 | 4 | |
| Jishan County, Yuncheng City, Shanxi Province | 33 | 33 | 12 | |
| Mian County, Hanzhong City, Shaanxi Province | 50 | 50 | 3 | |
| Fu County, Yan 'an City, Shaanxi Province | 11 | 11 | 3 | |

| | | | | |
|---|---|---|---|---|
| Ganquan County, Yan 'an City, Shaanxi Province | 4 | 3 | 2 | |
| Songjiang District, Shanghai | 1 | 1 | | 1 |
| Cangxi County, Guangyuan City, Sichuan Province | 60 | 58 | 3 | 2 |
| Chaotian District, Guangyuan City, Sichuan Province | 41 | 40 | 4 | 1 |
| Jizhou District, Tianjin | 25 | 25 | 6 | |
| Tengchong City, Baoshan City, Yunnan Province | 8 | 8 | | 2 |
| Anji City, Huzhou City, Zhejiang Province | 9 | 7 | 5 | |
| Dinghai District, Zhoushan City, Zhejiang Province | 5 | 5 | 5 | |
| Daishan County, Zhoushan City, Zhejiang Province | 82 | 25 | 1 | |
| Shengsi County, Zhoushan City, Zhejiang Province* | 31 | 31 | 1 | 1 |
| Kaizhou District, Chongqing | 6 | 3 | | 3 |

* parth- ticks collected around SFTS patients' house

### Counties located in Dabie Mountain

**Table S2** Location and polyploid information of Asian longhorned ticks (ALTs) samples from overseas.

| Location | Number of ALT DNA samples | Ploidy |
|---|---|---|
| Oita Prefecture, Japen | 3 | Bi- |
| Kagoshima Prefecture, Japan | 5 | Parth- |
| Jeju Province, Republic of Korea | 2 | Parth- |
| New Jersy, USA | 1 | Parth- |
| New South Wales, Austrilia | 2 | Parth- |
| Queensland, Austrilia | 2 | Parth- |

| | | |
|---|---|---|
| Okayama Prefecture, Japan | 5 | Parth- |
| South Island, New Zealand | 1 | Parth- |

**Appendix Figure.**

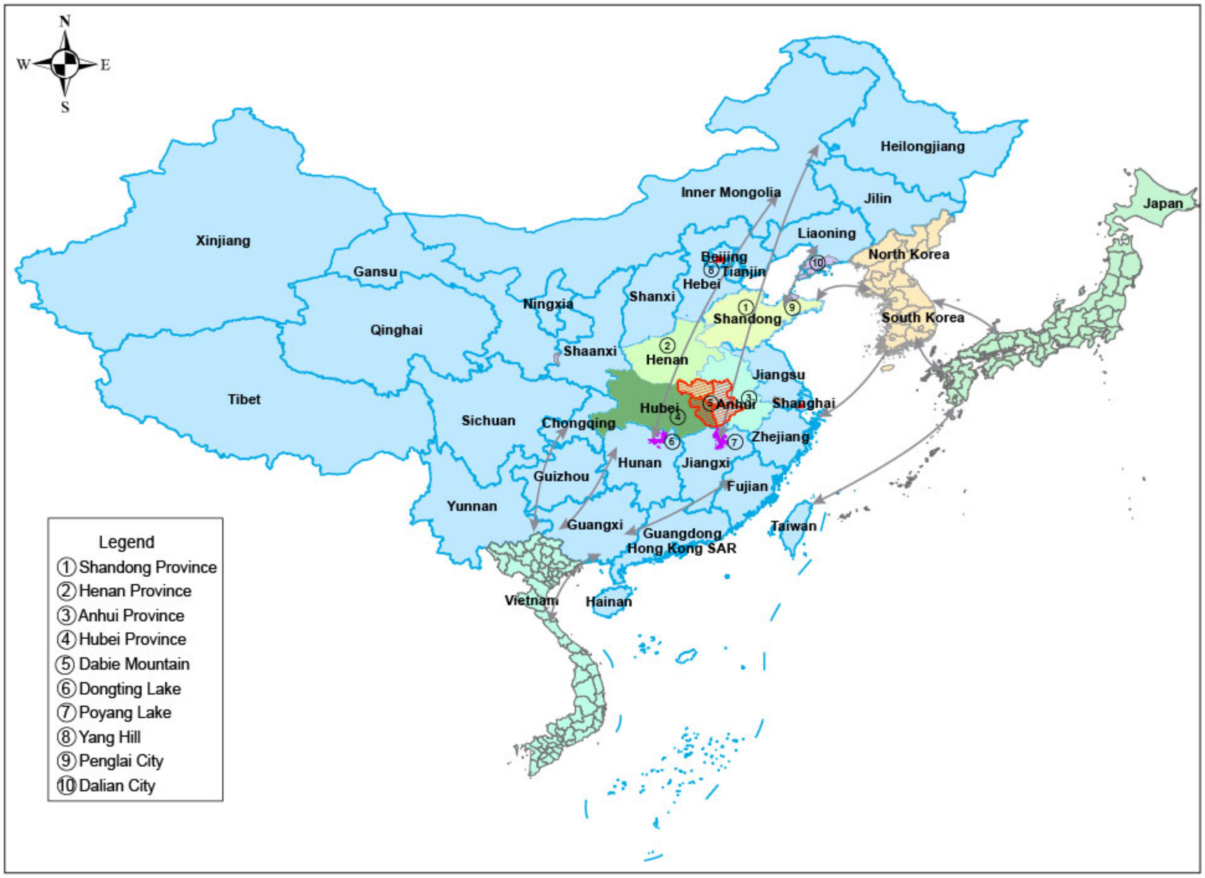


**Figure S1**. **Locations mentioned in this report. Arrows indicate bird migration routes.**

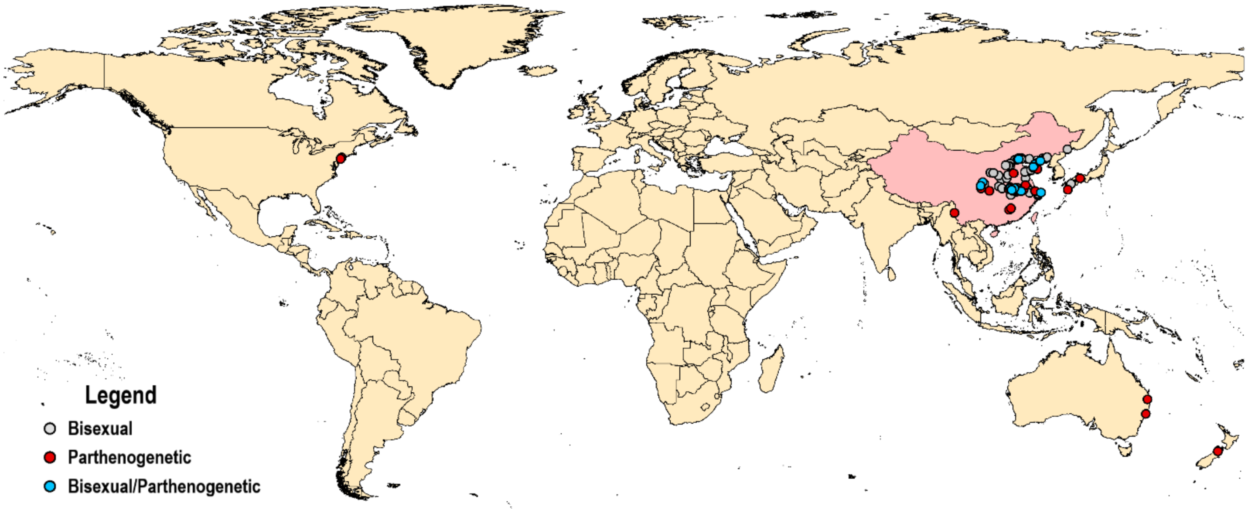


**Figure S2**. **Bi- and parth- Asian longhorned ticks collected in the Asia-Pacific area.** Red dots indicate parth-; gray dots indicate bi-; blue dots indicate both bi- and parth-.

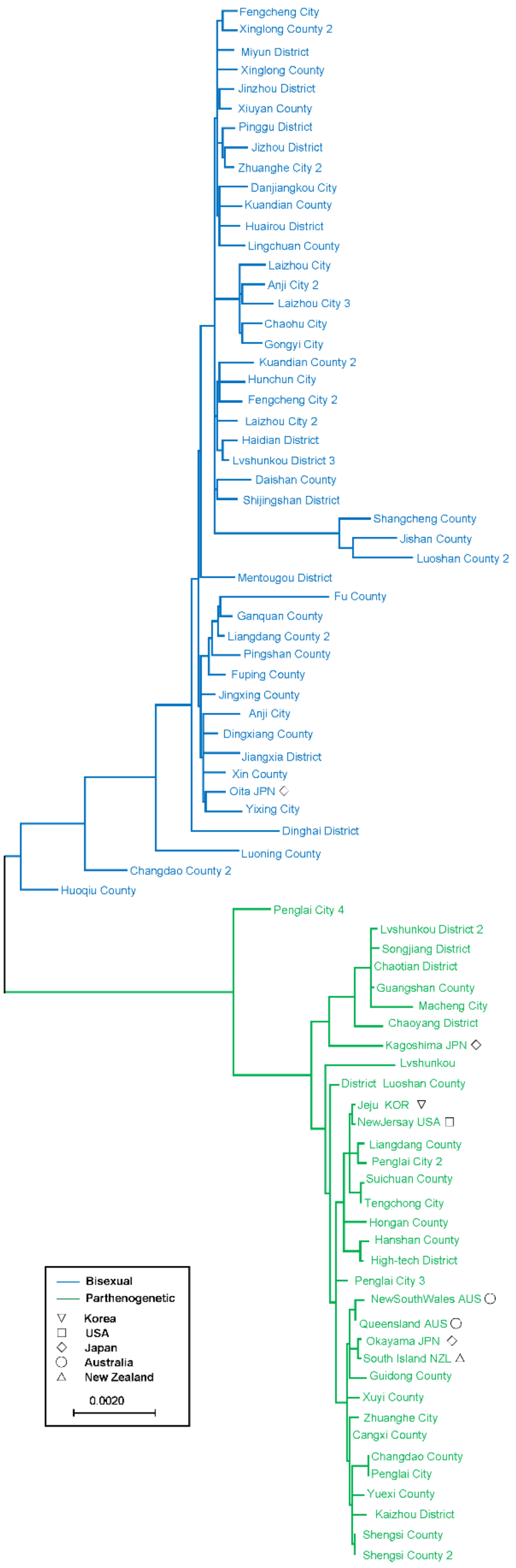
Fengcheng City
Xinglong County 2
Miyun District
Xinglong County
Jinzhou District
Xiuyan County
Pinggu District
Jizhou District
Zhuanghe City 2
Danjiangkou City
Kuandian County
Huairou District
Lingchuan County
Laizhou City
Anji City 2
Laizhou City 3
Chaohu City
Gongyi City
Kuandian County 2
Hunchun City
Fengcheng City 2
Laizhou City 2
Haidian District
Lvshunkou District 3
Daishan County
Shijingshan District
Shangcheng County
Jishan County
Luoshan County 2
Mentougou District
Fu County
Ganquan County
Liangdang County 2
Pingshan County
Fuping County
Jingxing County
Anji City
Dingxiang County
Jiangxia District
Xin County
Oita JPN
Yixing City
Dinghai District
Luoning County
Changdao County 2
Huoqiu County
Penglai City 4
Lvshunkou District 2
Songjiang District
Chaotian District
Guangshan County
Macheng City
Chaoyang District
Kagoshima JPN
Lvshunkou
District Luoshan County
Jeju KOR
NewJersay USA
Liangdang County
Penglai City 2
Suichuan County
Tengchong City
Hongan County
Hanshan County
High-tech District
Penglai City 3
NewSouthWales AUS
Queensland AUS
Okayama JPN
South Island NZL
Guidong County
Xuyi County
Zhuanghe City
Cangxi County
Changdao County
Penglai City
Yuexi County
Kaizhou District
Shengsi County
Shengsi County 2
Bisexual
Parthenogenetic
Korea
USA
Japan
Australia
New Zealand
0.0020

**Figure S3. Maximum likelihood tree established with the mitochondrial genomes of Asian longhorned ticks (ALTs) collected in the Asian-Pacific region.** Multiple ALTs from the same county were marked by number.

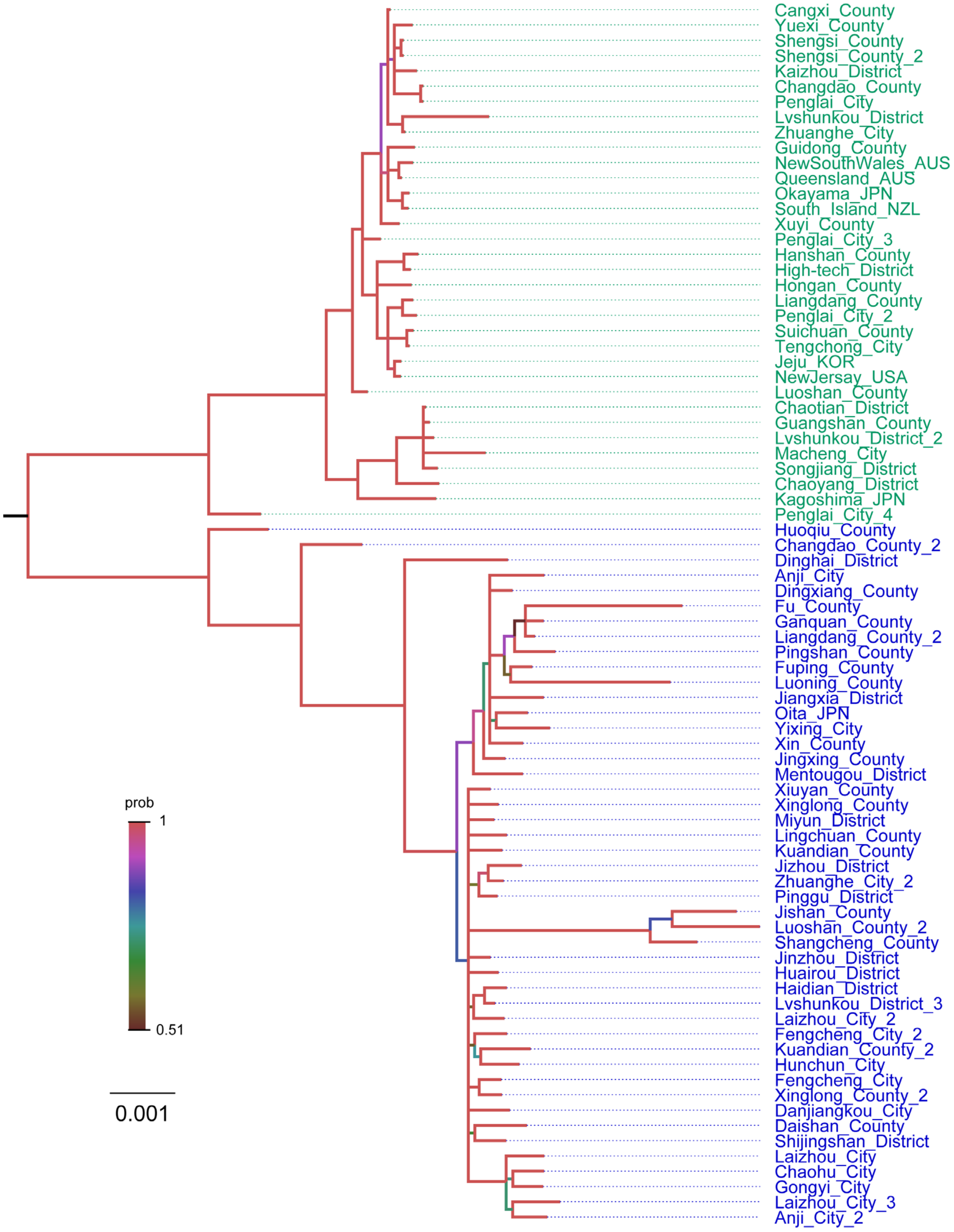

**Figure S4. Bayesian phylogenetic tree of parthenogenetic (green label) and Bisexual (blue label) Asian longhorned ticks (ALTs).** The color of the branches indicates the posterior probability. This tree was build by MrBays-3.2.7 (http://nbisweden.github.io/MrBayes/index.html). with 1,500,000 generations, and the average standard deviation of split frequencies of the tree is less than 0.01.

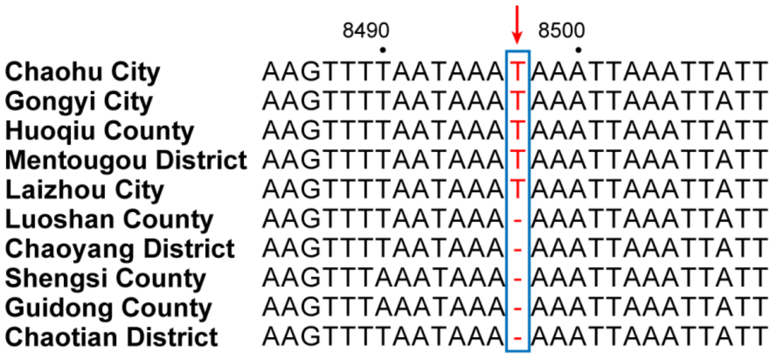


**Figure S5**. **Mitochondrial genome alignment between parthenogenetic and bisexual Asian longhorned ticks (ALTs) populations.** The upper five sequences were from bisexual ALTs and the lower five sequences were from parthenogenetic ALTs.

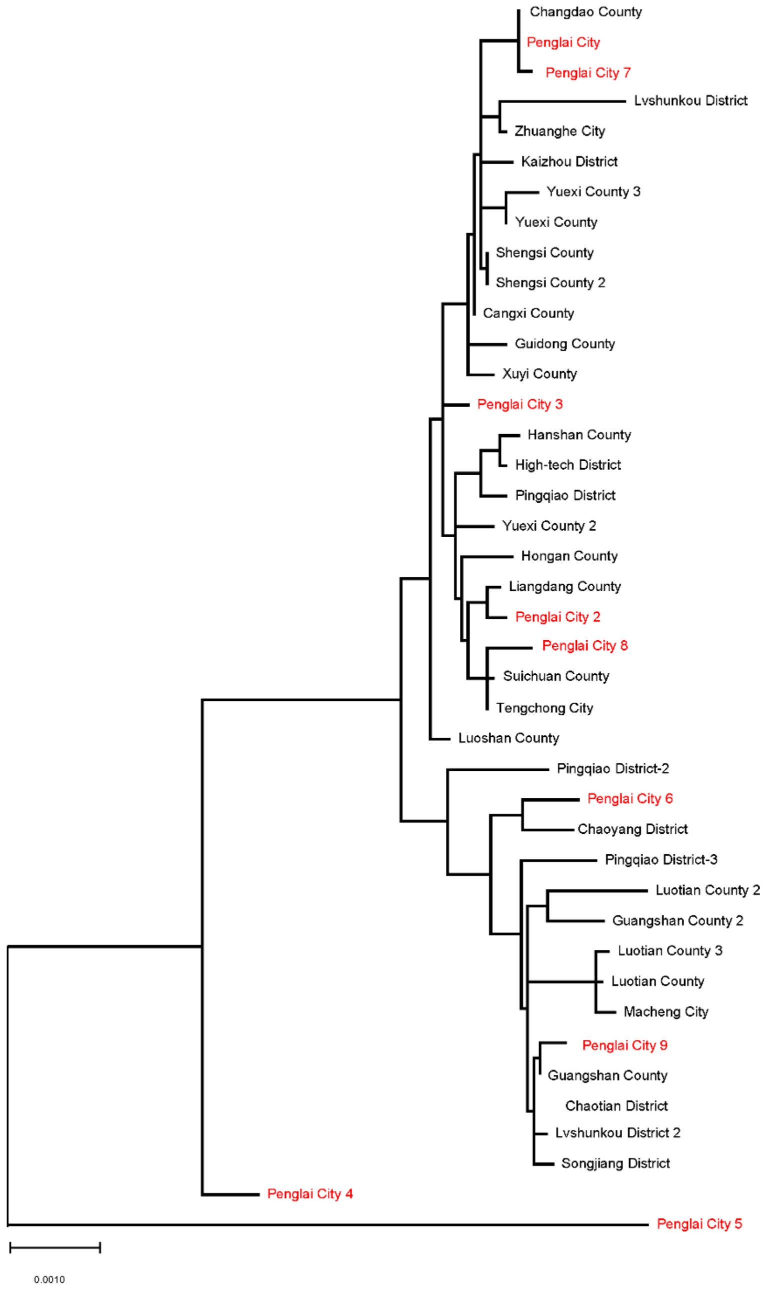


**Figure S6**. **Phylogenetic analysis of parthenogenetic ticks collected from Penglai City (red) and samples from 15 provinces in China.** Maximum likelihood tree established with the mitochondrial genomes of ticks. Multiple ticks from the same county were marked by number.
